# Effects of cochlear synaptopathy on spontaneous and sound-evoked activity in the mouse inferior colliculus

**DOI:** 10.1101/381087

**Authors:** Luke A. Shaheen, M. Charles Liberman

**Affiliations:** Oregon Hearing Research Center, Oregon Health &Science University, Portland OR, USA; Department of Otology and Laryngology, Harvard Medical School, Boston MA, USA; Eaton-Peabody Laboratories, Massachusetts Eye &Ear Infirmary, Boston MA, USA

## Abstract

Tinnitus and hyperacusis are life-disrupting perceptual abnormalities that are often preceded by acoustic overexposure. Animal models of overexposure have suggested a link between these phenomena and neural hyperactivity, i.e. elevated spontaneous rates (SRs) and sound-evoked responses. Prior work has focused on changes in central auditory responses, with less attention paid to the exact nature of the associated peripheral damage. The demonstration that acoustic overexposure can cause cochlear nerve damage without permanent threshold elevation suggests this type of peripheral damage may be a key elicitor of tinnitus and hyperacusis in humans with normal audiograms. We addressed this idea by recording responses in the mouse inferior colliculus (IC) following a bilateral, neuropathic noise exposure. Two wks post-exposure, mean SRs were unchanged in mice recorded while awake, or under anesthesia. SRs were also unaffected by more intense, or unilateral exposures. These results suggest that neither neuropathy nor hair cell loss are sufficient to raise SRs in the IC, at least in mice. However, it’s not clear whether our mice had tinnitus. Tone-evoked rate-level functions at the CF were steeper following exposure, specifically in the region of maximal neuropathy. Furthermore, suppression driven by off-CF tones and by ipsilateral noise were also reduced. Both changes were especially pronounced in neurons of awake mice. These findings align with prior reports of elevated acoustic startle in neuropathic mice, and indicate that neuropathy may initiate a compensatory response in the central auditory system leading to the genesis of hyperacusis.

## INTRODUCTION

Tinnitus, the phantom perception of sound, is often preceded by an episode of acoustic overexposure (Eggermont and Roberts, 2004). Since the anomalous percept occurs in the absence of external sound, the search for the neurophysiological basis of tinnitus has long focused on patterns and rates of spontaneous discharge in the auditory pathway. The auditory nerve, the primary sensory neurons, normally show a range of spontaneous rates (SRs) from near 0 to over 100 sp/sec (Liberman, 1978; Taberner and Liberman, 2005). Curiously, acoustic overexposure, in animal models, almost always decreases or eliminates spontaneous discharge, whereas it often causes increased SRs across the central auditory pathway, including in the ventral and dorsal cochlear nucleus (DCN), inferior colliculus (IC) and auditory cortex (Brozoski et al., 2002; Bauer et al., 2008; Seki and Eggermont, 2003). Since overexposed animals have been inferred to have tinnitus-like percepts by a variety of behavioral tests, hyperactivity is hypothesized to be a neural correlate of tinnitus (Brozoski et al., 2002; Longenecker and Galazyuk, 2011).

However, the relationship between neuronal hyperactivity and a tinnitus-like percept is not straightforward: in animals given identical noise exposures, not all demonstrate changes in the behavioral assays indicating a tinnitus like-percept, such as a reduction in pre-pulse inhibition (PPI) of the startle reflex induced by a preceding gap in an otherwise continuous noise background. While SRs in the DCN are elevated only in animals with reduced gap-induced PPI (Dehmel et al., 2012a; Koehler and Shore, 2013; Li et al., 2013), SRs in the IC and auditory cortex are elevated regardless of PPI (Engineer et al., 2011; Coomber et al., 2014; Ropp et al., 2014). Furthermore, DCN ablation before noise exposure prevents the emergence of a tinnitus-like percept as assessed by conditioned suppression, whereas the same ablation performed 3-5 months after noise exposure has no effect (Brozoski and Bauer, 2005; Brozoski et al., 2012). Hyperactivity in the IC follows a similar pattern: cochlear ablation up to 4 wks post-exposure causes a return of SRs to control levels, but ablation ≥ 8 wks post-exposure does not (Mulders and Robertson, 2009, 2011).

It is not clear how much cochlear damage, if any, is necessary to cause hyperactivity in central auditory pathways. Although threshold shift is a primary risk factor for tinnitus in humans (Roberts et al., 2010), tinnitus can occur without threshold shifts (Gu et al., 2010; Schaette and McAlpine, 2011; Gu et al., 2012). Elevated SRs and behavioral evidence for tinnitus have also been observed in animals where acoustic overexposure caused little to no permanent threshold shift (Bauer et al., 2008; Koehler and Shore, 2013). Moderate noise exposures that do not cause hair cell damage or permanent threshold shift (PTS) can cause a loss of synapses between cochlear hair cells and auditory nerve terminals, followed by a slow degeneration of auditory-nerve cell bodies and central axons (Kujawa and Liberman, 2009). Such a cochlear neuropathy would not be apparent in an audiogram, but may be detected by reductions in evoked response amplitudes, such as the auditory brainstem response (ABR) or envelope following response (EFR: Bharadwaj et al., 2015; Shaheen et al., 2015). Since tinnitus patients with normal thresholds have reduced ABR wave I amplitudes (Gu et al., 2012), it is possible that these individuals also have cochlear neuropathy.

Hyperacusis, an intolerance to moderate- or high-level sounds, often co-occurs with tinnitus and like tinnitus, is often preceded by an episode of acoustic overexposure (Kreuzer et al., 2012). In humans with hyperacusis, sound-evoked fMRI responses are increased in the auditory midbrain, thalamus, and cortex (Gu et al., 2010). In animal models, acoustic overexposure can result in elevated sound-evoked responses in the ascending auditory pathways (Salvi et al., 1990; Cai et al., 2009; Auerbach et al., 2014), and hypersensitivity to loud sounds (Hickox and Liberman, 2014; Manohar et al., 2017). Since hyperacusis can occur in humans with normal audiograms, but is often associated with acoustic overexposure, cochlear neuropathy may also be a key catalyst in the generation of hyperacusis.

Most prior work on noise-induced central hyperactivity did not quantify the condition of hair cells and/or cochlear nerve fibers, nor the possible correlations between peripheral histopathology and central pathophysiology (but see Hesse et al., 2016; Asokan et al., 2018). While identically exposed animals develop behavioral evidence of tinnitus at rates ranging from 30 to 70% (Engineer et al., 2011; Dehmel et al., 2012b; Li et al., 2013; Rüttiger et al., 2013), these studies did not ask whether variability was related to differences in peripheral insult. Here, our aim was to test the idea that both spontaneous and sound-evoked hyperactivity are caused by cochlear neuropathy *per se*, by measuring responses in the IC of mice exposed to noise calibrated to induce cochlear neuropathy with minimal hair cell loss, and to more intense noise that caused both neuropathy and hair cell loss. Spontaneous rates were not increased in either group relative to controls, indicating that neither cochlear neuropathy nor hair cell loss is sufficient to cause increased SRs in the mouse IC. However, tone-evoked rate-level functions were steeper following exposure, specifically in the region of maximal cochlear neuropathy, suggesting that neuropathy plays a role in hyperacusis.

## METHODS

### Animals and groups

Seven wk-old male CBA/CaJ mice were exposed to octave-band noise (8-16 kHz) for 2 hrs. Noise calibration to target SPL was performed immediately before each exposure session. Control mice were of the same age, gender, and strain, but were not exposed to the noise. Physiological experiments were always performed 1-3 wks after the noise exposure. When anesthesia was used for noise exposure, it was a ketamine/xylazine mix (100/20 mg/kg, respectively, i.p) with booster injections as needed. All procedures were approved by the Institutional Animal Care and Use Committee of the Massachusetts Eye and Ear Infirmary.

Two experimental groups were exposed awake and unrestrained to the 2-hr noise band at 98 dB SPL, a level/duration known to produce “pure” cochlear synaptopathy in both ears, i.e. loss of cochlear synapses without any hair cell loss or permanent threshold shift (Kujawa and Liberman, 2015; Shaheen et al., 2015; Suzuki et al., 2016; Valero et al., 2016). IC recordings were then made under either awake (n=6) or anesthetized (n=10) conditions, and results were compared to similar recordings in unexposed controls under either awake (n=8) or anesthetized (n=13) conditions. After recordings, a subset of exposed and control animals was sacrificed for histopathological analysis, to confirm the synaptopathy phenotype that has been replicated in many other studies from our group (Kujawa and Liberman, 2015; Shaheen et al., 2015; Suzuki et al., 2016; Valero et al., 2016). A third experimental group (n=6) was exposed awake and unrestrained to the noise band at 103 dB SPL, a level/duration designed to cause synaptopathy plus significant hair cell damage and permanent threshold shifts. IC recordings were made from this group under awake conditions (n=6), but none of these ears was processed histologically.

To evaluate possible differences between bilateral and unilateral noise damage, an additional three groups were exposed unilaterally while under ketamine/xylazine anesthesia using a small tweeter coupled to the ear canal via a speculum. Exposures were conducted in a warm sound-proofed room (30° C), and a stable anesthetic plane was maintained with booster injections as needed. To minimize contralateral exposure, mice were placed on their side with saline-soaked cotton in the contralateral ear canal. The three groups were exposed to the 2-hr noise band at either 101 dB (n=2), 103 dB (n=2), or 104 dB SPL (n=2) and then used for IC recordings under anesthetized conditions. None of these ears was processed histologically.

### Preparation for IC recordings in anesthetized mice

A single session was conducted in each animal, occurring 1-3 wks post-exposure. Mice were anesthetized with a ketamine/xylazine mix (100/20 mg/kg, respectively, i.p) with booster injections as needed. Surgical procedures and recordings were conducted in an acoustically and electrically shielded room at 30° C. Prior to surgery, DPOAEs and ABRs were measured as described below. Then, the pinnae were removed bilaterally to allow for closed-field acoustic stimulation using two custom acoustic assemblies, each containing two electrostatic drivers (CUI CDMG15008-03A) and an electret condenser microphone (Knowles FG-23329-P07). The assemblies were calibrated with a ¼-inch Bruel and Kjaer condenser microphone and in-ear calibrations were performed at the onset of each experiment.

A small craniotomy was made over the central nucleus of the IC using a scalpel. The dura mater was left intact and covered with high-viscosity silicon oil. Recordings were made with a 16-channel, single-shank silicon probe (25 or 50 μm contact separation, 177 μm^2^ contact area; NeuroNexus Technologies). Electrodes were inserted dorso-ventrally, along the tonotopic axis, and advanced using a micropositioner (Kopf 607-C). While the central nucleus of the IC is easy to visualize in mice, electrode placement was confirmed by the tonotopic organization and sharp frequency tuning.

### Preparation for IC recording in awake mice

Two to five days following noise exposure, mice were anesthetized with ketamine and xylazine. The skin overlying the frontal and parietal portions of the skull was retracted, and a titanium headplate was glued to the skull using acrylic bonding material to allow for head-fixed recordings (C&B MetaBond, Parkell). Buprenorphine (0.1 mg/kg s.c.) and meloxicam (0.03 mg/kg s.c.) were given, and animals were allowed to recover for at least 48 hours. Recording sessions began as soon as one wk post-exposure and continued until three wks post-exposure; each session lasted up to three hours. One to nine (median 3) recording sessions were conducted in each mouse. At the onset of each session, mice were anesthetized with isoflurane in oxygen (2%, 2 L/min). In the first session, a craniotomy was made over the IC as in the anesthetized recordings. A ring around the craniotomy site was formed using a UV-cure resin, and the craniotomy was filled with high-viscosity silicon oil. The same craniotomy was used in subsequent sessions. Cyanoacrylate was used to glue a 1.8 mm rigid plastic tube in each ear canal. Isoflurane was discontinued, and the mouse was transferred to the recording apparatus, where it was head-fixed but free to run on a 22-cm diameter turntable beneath its feet. The ear tubes were coupled to the acoustic assemblies for closed-field sound delivery, and in-ear calibration was performed. DPOAEs were monitored throughout the session to ensure that the acoustic delivery remained constant. Animals were continuously monitored, and the session was ended for the day if they showed signs of persistent distress. Recordings were made using the same 16-channel Neuronexus probes as for experiments on anesthetized mice. At the end of each session, the mice were briefly anesthetized with isoflurane, and the ear tubes were removed. The craniotomy was cleaned and flushed with saline, then covered with a thin layer of bacitracin and sealed with UV-cure resin. 3-wks post exposure, animals were anesthetized with ketamine/xylazine and DPOAEs and ABRs were recorded. At the conclusion of these recordings, mice were sacrificed for histology.

### IC recording

Custom LabVIEW and MATLAB software controlling National Instruments 24-bit digital input/output boards generated all stimuli at a 200 kHz sampling rate. Raw signals from the electrodes were digitized at 32 bits, 24.4 kHz (RZ5 BioAmp Processor; Tucker-Davis Technologies) and stored in binary format. For awake recording, the point-by-point median of all channels was subtracted from each channel, greatly reducing motion-induced artifacts. Single units were isolated online using a sorting method based on principle-components analysis (PCA) in SpikePac software (Tucker-Davis Technologies). Units were later reprocessed offline using a combination of custom MATLAB software and the Ultra Mega Sort 2000 package (Hill et al., 2011): raw waveforms were filtered both forward and backward using a fourth-order 300 – 5000 Hz Butterworth filter. Spikes were detected by threshold crossings above 3.5 standard deviations of the waveform (measured over intervals far removed from sound presentation). Threshold crossings occurring < 0.6 ms after another spike were removed to prevent double counting due to noise. Single units were then sorted by: aligning spike waveforms to the peak, reducing dimensionality using PCA, overclustering using k-means (k>32), and merging clusters based on the interface energy between clusters and violations of the absolute refractory period (Hill et al., 2011). Candidate single units were then checked for good isolation by their separation from other clusters, estimated number of sub-threshold (missed) spikes, and interspike interval distributions. The majority of channels contained 0 or 1 well-isolated single units, but some channels contained up to 2 single units. In some cases, especially for awake recordings, movement of the brain relative to the skull caused single units to drift from channel to channel, at times back and forth between channels. In these cases spikes were detected and aligned from included channels independently, waveforms were concatenated, and the single-channel procedure was followed beginning with PCA.

### Multi-unit Activity

During most recordings, only 2-4 channels contained single-units. We processed the remaining multiunit activity in two ways: 1) thresholded multi-unit activity (tMUA) was computed by setting the spike threshold to 3.5 standard deviations of the neural activity (filtered 300 - 5000 Hz) during periods of silence presented during recording of frequency response areas (FRAs), and 2) the envelope of multi-unit activity (eMUA) was computed by band-pass filtering from 300 to 5000 Hz, computing the absolute value, and then computing the average across all repetitions of the stimulus (Supèr and Roelfsema, 2005; Kayser et al., 2007; Choi et al., 2010). In both cases, sites that had strong single-unit responses were excluded to avoid biasing the sample.

### Stimuli

The search stimulus was a broadband noise burst presented at 60 dB SPL, alternating sides on each presentation.

### Frequency Response Areas

FRAs were collected by pseudorandom presentation of tone pips (50 ms on, 200 ms off, 4 ms raised cosine onset/offset ramps) to the ear contralateral to the recording site. Between 2 and 4 trials were collected per tone/level combination. Tone frequencies were of 3 kHz to 70 kHz, spaced in 0.1 octave increments. The specific subset was set to cover the response range of the neurons under investigation (generally 2 to 3 octaves wide). In mice recorded under anesthesia, levels were 10 to 80 dB SPL in 5 dB steps, in mice recorded awake, levels were −5 to 80 dB in 5 dB steps. For both groups, a silent condition was added to the list of levels to be randomized. Spontaneous rate (SR) was calculated over the entire 250 ms of this these silent trials, averaged across trials and ‘frequencies.’ Thus, the total time used to calculate SR ranged from 10 to 30 seconds. Discharge rates in response to tones were calculated over a time window 5 to 55 ms *re* stimulus onset, and used to construct the FRA display matrices. The excitatory region of the FRA was determined by 1) measuring the mean and standard deviation of the SR, 2) zeroing all FRA pixels < 5 standard deviations above mean SR, grouping non-zero matrix cells into regions connected by edges, and finding the region with the largest average rate * (number of cells)^2^. This procedure successfully identified the excitatory region for the largest range of FRA shapes, and manual intervention was rarely required. Within the excitatory region, the characteristic frequency (CF) was defined as the frequency with the highest response at the lowest stimulus level, and the best frequency (BF) as the frequency producing the largest average response across all levels. The bandwidth 10 dB above threshold (BW_10_) was used to measure the sharpness of excitatory tuning.

### Binaural-Noise Response Areas

Binaural-noise response areas were collected by pseudorandom presentation of simultaneous, dichotic noise bursts (50 ms on, 200 ms off, 4 ms raised cosine onset/offset ramps). Noise was limited from 4 kHz to 80 kHz and shaped to be flat-spectrum by correction for the frequency response of the acoustic system. Ipsilateral and contralateral levels were randomly and independently varied, and 5 trials were collected per level combination. Levels were chosen to encompass the dynamic range of the neurons under investigation, in anesthetized mice this was generally 20 to 70 dB SPL, while in awake mice this was 10 to 70 dB SPL, both in 5 dB steps. As for the FRAs, a silent condition was added to the list of levels to be randomized, so that responses to both contralateral noise alone and ipsilateral noise alone were included in the stimulus set. Rate was averaged over a time window 5 to 55 ms *re* stimulus onset.

### Fits to rate-level functions

Rate-level functions at CF and BF were calculated from FRAs and fit with a four-parameter model (Sachs and Abbas, 1974; Heil et al., 2011). This model specifies firing rate *r* as a function of sound pressure *P* (in Pa) by:

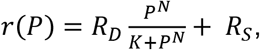

and

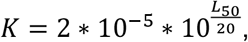

where R_D_ is the driven rate, R_S_ is the spontaneous rate, N characterizes the steepness of the curve, and L_50_ gives the level at 50% of the driven rate range. Non-monotonic functions were manually selected and fit using the sum of two such models, yielding seven parameters:

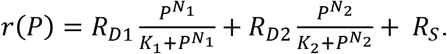

The slope of the rising portion of the function was used in population comparisons. Curves were fit by *fminsearchbnd*, a bounded version of the built-in MATLAB function *fminsearch* using a least-squares method. The fit was then refined by a robust procedure that iteratively refined weights of each data point before re-fitting.

### DPOAEs and ABRs

For measurement of ABRs and DPOAEs, stimuli were presented unilaterally, with the mouse on its side, and the acoustic assembly just above the ear canal. After data were collected for one ear, the procedure was repeated for the other ear.

ABRs were recorded differentially from subdermal needle electrodes with the common ground at the base of the tail. A dorsal-ventral incision was made in the pinna at notch between the tragus and antitragus to allow direct visualization of the eardrum. For mice that had undergone IC recordings under anesthesia, the electrode montage was vertex (positive) to ipsilateral pinna, with the latter just caudal to the intertragal notch (see Shaheen et al 2015). For those that had undergone awake recordings, the headplate prohibited placing the vertex electrode. Therefore, active electrodes were placed either 1) headplate to ipsilateral pinna, or 2) contralateral pinna to ipsilateral pinna. ABR amplitudes were more variable for configuration 1); thus, results are reported for configuration 2). Pinna electrodes were placed slightly more ventral along the antitragus in order to increase the early ABR waves (pinna_V_ in Shaheen et al 2015).

Responses were amplified 10,000X using two Grass P511 amplifiers with a 0.3-3 kHz passband. ABRs were evoked with 5-msec tone-pips with 0.5 msec cos^2^ rise—fall, presented in alternating polarity at a rate of 40/s. Tone-pip frequencies were 11.3 or 32 kHz. Trials where the response amplitude exceeded 15 μV were rejected; 512 artifact-free trials of each polarity were averaged to compute ABR waveforms. Threshold was defined by visual inspection of the stacked waveforms as the lowest level at which a reproducible peak or trough appears, which usually occurs one level-step below that at which peak-to-peak amplitude begins to grow.

DPOAEs were recorded in the ear canal sound pressure in response to two tones, f_1_ and f_2_, each presented to separate speakers to reduce system distortion (frequency ratio f_2_/f_1_ = 1.2, and level difference L_1_ = L_2_ + 10 dB). DPOAE response was measured at 2f_1_-f_2_ by Fourier analysis of the ear-canal sound pressure waveform. Stimulus duration was 1.6 seconds at each level combination (L_2_ varied from 20 to 80 dB SPL in 5 dB steps). Threshold was defined as the interpolated f_2_ level producing a DPOAE of 5 dB SPL.

### Cochlear Immunostaining and Innervation Analysis

Mice were perfused intracardially with 4% paraformaldehyde. Cochleas were decalcified, dissected into half-turns and incubated in primary antibodies: 1) mouse (IgG1) anti-CtBP2 from BD Biosciences at 1:200 and 2) mouse (IgG2) anti-GluA2 from Millipore at 1:2000. Primary incubations were followed by 60-min incubations in species-appropriate secondary antibodies. Cochlear lengths were obtained for each case, and a cochlear frequency map computed using a custom ImageJ plugin (http://www.masseyeandear.org/research/otolaryngology/investigators/laboratories/eaton-peabody-laboratories/epl-histology-resources/) that translates cochlear position into frequency according to the published map for the mouse (Müller et al., 2005; Taberner and Liberman, 2005). Confocal z-stacks from each ear were obtained using a 63x glycerol-immersion objective (N.A.=1.3) at 3.17X digital zoom on a Leica TCS SP5 confocal. Synapses in the IHC area were counted using Amira (Visage Imaging) to find the xyz coordinates of all the ribbons (CtBP2-positive puncta), and custom re-projection software was then used to assess the fraction of ribbons with closely apposed glutamate-receptor patches (GluA2 puncta).

### Statistical Analysis

Statistical testing was performed in MATLAB; functions used are indicated in italics. For spontaneous rates, distributions were normal (assessed with Lilliefors test), so differences were assessed using a 3-way ANOVA (*anovan*). Significant post-hoc differences were assessed using *multcompare* with a Tukey-Kramer correction for multiple comparisons. Threshold, maximum driven rates, and slopes were not normally distributed, especially for multi-units. Therefore, differences were assessed with paired rank-sum tests. 6 comparisons were made to test effects of exposure, and a Bonferroni-Holm correction was applied. To test effects of anesthesia, populations were pooled across exposure for each frequency bin. Therefore, these tests were corrected for 3 comparisons.

## RESULTS

### Noise-induced synaptic loss after reversible noise-Induced threshold shift

We exposed two groups of 7-wk old mice to an 8-16 kHz, 98 dB SPL noise band for 2 hrs, which reliably causes substantial cochlear synaptopathy with minimal hair cell damage (Shaheen et al., 2015; Valero et al., 2016). One group was then used for awake IC recordings, and one for anesthetized IC recordings. For efficiency, the time-consuming histological analysis was completed only for the awake-recording group, since this exposure has produced such highly stereotyped results in many prior studies from the Liberman lab (Shaheen et al., 2015; Valero et al., 2016), thus only basic reconfirmation is necessary to ensure that some idiosyncratic difference in exposure technique has not altered the basic damage pattern.

As in prior work, this exposure resulted in a 40-50% loss of synapses between inner hair cells and auditory-nerve fibers in the basal half of the cochlea (**Fig 1A**). The effect of exposure was significant as assessed by a repeated-measures ANOVA (F_1,12_ = 101, p<<0.001). As expected, no damage to inner hair cells was observed (data not shown), and loss of outer hair cells was confined to the basal end of the cochlea (**Fig 1D**). Consistent with the histopathology, DPOAE thresholds were normal except at 45 kHz, where they were elevated by 5-10 dB in both exposed groups (**Fig 1F**). DPOAE amplitudes were also reduced at 45 kHz (**Fig 1E**), consistent with the DPOAE threshold shift, and with the small but significant OHC damage in that, and more basal, regions (**Fig 1D**). ABR thresholds were normal at both 11.3 and 32 kHz (**Fig 1C**), as expected from the DPOAE thresholds. Exposure significantly reduced ABR wave I amplitudes in both anesthetized- and awake-recording groups (F_1,4_ = 8.8, p=0.04; F_1,21_ = 9.4, p=0.006 respectively). In the anesthetized-recording group, post-hoc tests indicated ABR wave I amplitudes were significantly reduced at 32 kHz, but not at 11.3 kHz (red symbols in **Fig 1B**), consistent with the distribution of synaptic loss (**Fig 1A**). ABR amplitudes were significantly reduced at both frequencies in the awake-recording group, although ABR amplitudes in this group (green symbols in **Fig 1B**) are hard to interpret because the head plate prevented normal placement of electrodes (see Methods).

**Figure 1.**
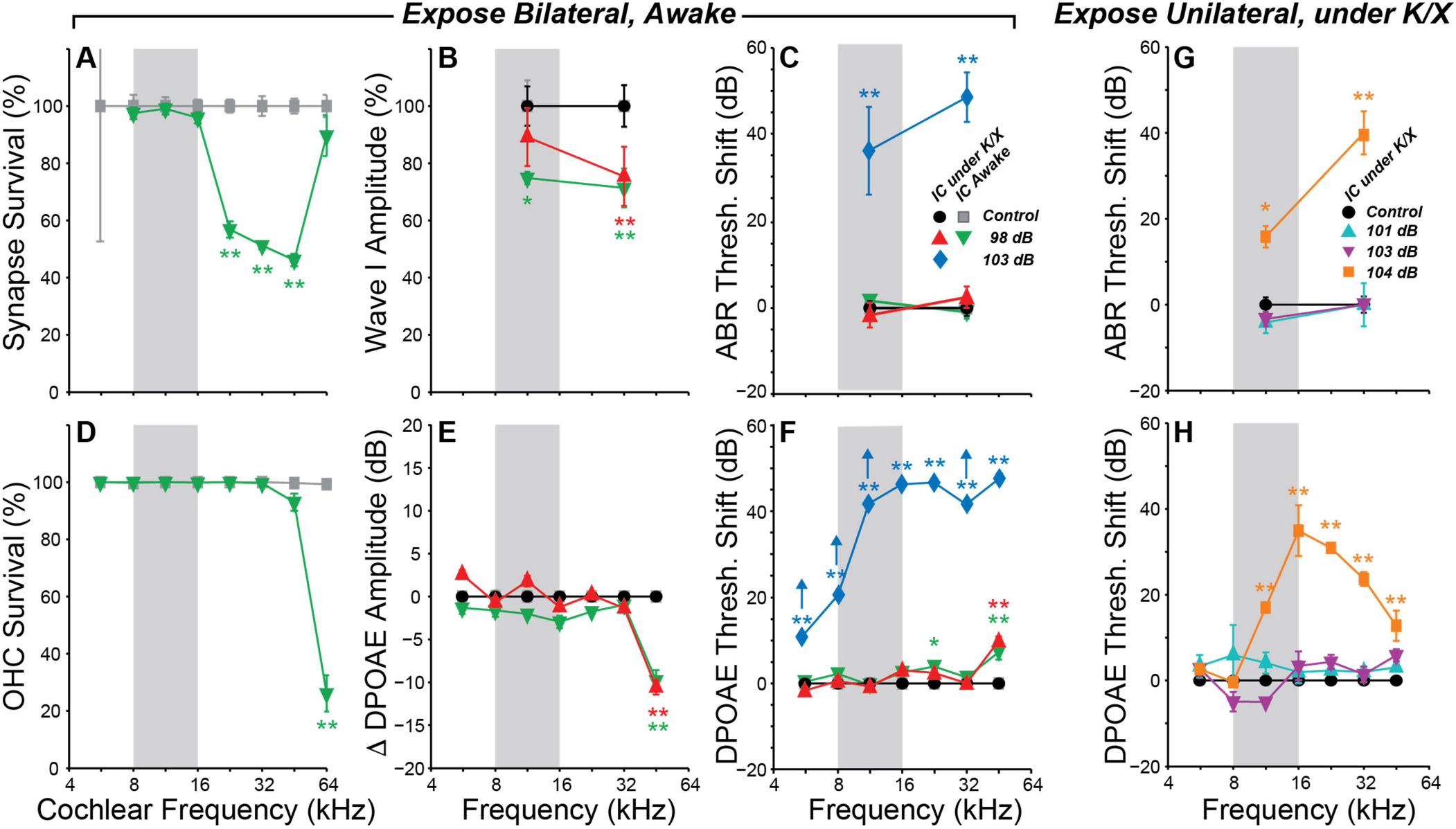
Summary histopathology and pathophysiology for all noise-exposure groups in the present study. Mice were awake for bilateral exposures (**A-F**) and under ketamine/xylazine (K/X) anesthesia for the unilateral exposures (**G-H**). **A,D**: Mean IHC synapse and OHC survival following 98 dB exposure in mice used for awake IC recordings. **B,E**: Mean change in ABR Wave I (**B**) and DPOAE (**E**) amplitudes, averaged over stimulus levels from 60 to 80 dB SPL. **C,F**: Mean ABR (**C**) and DPOAE (**F**) threshold shifts in mice used for IC recordings in either awake or anesthetized animals. Up arrows in **F** indicate that at least on animal showed no response at the highest stimulus level. Each group is normalized to its respective control. Key in C applies to all panels **A - F. G,H:** Mean ABR (**G**) and DPOAE (**H**) threshold shifts following unilateral exposures. Key in **G** also applies to **H**. Error bars in all panels indicate SEMs. Asterisks indicate significant effects of exposure (p<0.05 (single), p<0.001 (double) by Tukey-Kramer-corrected post-hoc test following a repeated-measures ANOVA.

Based on the highly reproducible patterns of hair cell and neural damage, we divided the cochlear frequency space into three regions: 1) ‘non-neuropathic’ (< 16 kHz), where noise exposure did not permanently alter thresholds, hair cells or synaptic counts, 2) ‘neuropathic’ (16-32 kHz), where exposure caused cochlear neuropathy without hair cell damage or threshold shift, and 3) ‘threshold-shift’ (>32 kHz), where exposure caused neuropathy with some hair cell damage and permanent threshold shift.

### Spontaneous rates in IC were unaffected by bilateral neuropathic noise exposure

We examined the central nucleus of the inferior colliculus (IC) of noise-exposed mice for signs of hyperactivity via single-unit responses using single-shank silicon electrodes. In the first group of mice, responses were measured under ketamine/xylazine anesthesia, in a single session 1-3 wks following noise exposure. SRs ranged from 0-50 spikes/sec in both control and noise-exposed mice, but were not affected by noise-exposure in any range of characteristic frequency (CF) (**Fig 2A**).

**Figure 2.**
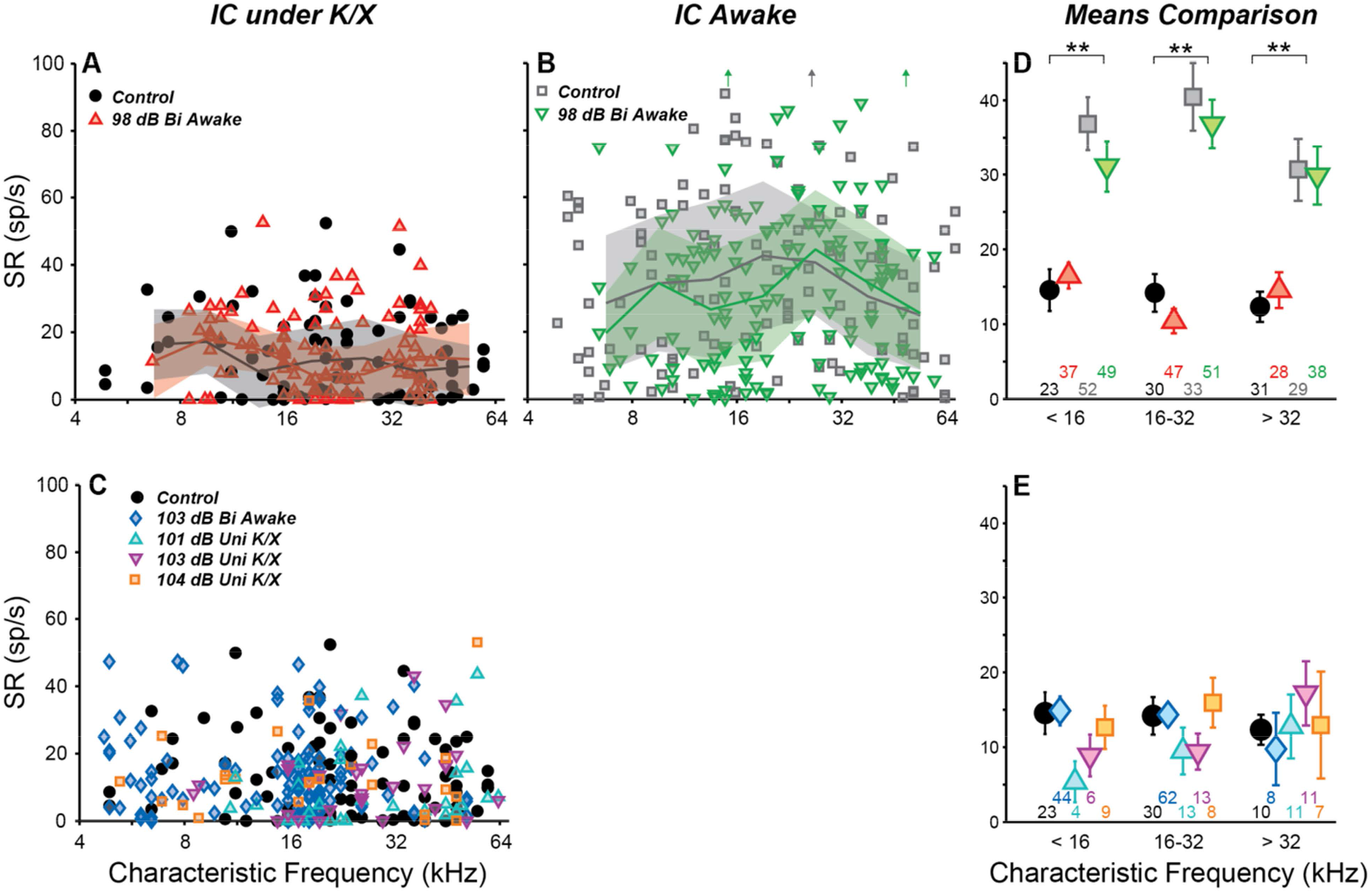
Noise exposure does not affect spontaneous rates (SRs) in IC. **A,B,C:** SRs from isolated IC neurons in control vs. exposed mice, recorded in anesthetized (**A,C**) or awake (**B**) preparations. Lines and shading in **A** and **B** indicate medians and interquartile ranges, pooled over one-octave bands. Off-axis SRs (up arrows in **B**) were 110, 131, and 104 sp/s from left to right. **D,E:** Mean SRs binned into three CF regions, as indicated. Black asterisks indicate significant effects of anesthesia (p<0.001, Tukey-Kramer-corrected post-hoc test following a 3-way ANOVA). Numbers below points indicate the number of single units contributing to each average. Keys in **A** and **B** also apply to **D**; key in **C** applies to **D**.

Anesthesia has a strong effect on IC responses, including SRs (Torterolo et al., 2002; Chung et al., 2014). Therefore, we recorded responses from a group of awake, head-fixed mice. One to ten (median three) recording sessions were conducted in each mouse, starting from 1 wk post-exposure and ending 3 wks post-exposure. SRs ranged from 0-130 spikes/sec and were significantly elevated relative to anesthetized mice (**Fig 2B**). However, there was no effect of noise exposure on SRs.

SR means were calculated over three CF regions and the significance of effects was evaluated using a three-way ANOVA (**Fig 2D**). The main effect of anesthesia was highly significant (F_1,436_ = 110, p<<0.001), but effects of exposure and frequency region were not (F_1,436_ = 0.9, p=0.3; F_2,436_ = 2.6, p=0.08). Post-hoc tests did not reveal significant effects of exposure in any frequency region.

### Spontaneous rates in IC were also unaffected by unilateral or more intense exposures

Most studies of noise-induced SR elevation in the IC have used more intense noise exposures, causing a greater amount of cochlear pathology (ex. Mulders and Robertson, 2009). Therefore, we raised the noise exposure level to 103 dB SPL while holding all other parameters constant. At this intensity, exposure caused > 40 dB of permanent threshold shift across a wide range of frequencies (**Fig 1C,F**, blue diamonds). Surprisingly, SR still remained unchanged relative to controls (**Fig 2E**).

While elevated SRs have been observed following bilateral exposures in a number of prior studies (Ma et al., 2006; Manzoor et al., 2012, 2013; Hesse et al., 2016), the majority of studies reporting SR increases have used unilateral exposures (Bauer et al., 2008; Mulders and Robertson, 2009; Longenecker and Galazyuk, 2011; Coomber et al., 2014; Ropp et al., 2014; Vogler et al., 2014). Thus, we set out to measure SRs in three groups of unilaterally exposed mice. The exposure band and duration were held constant, but mice were exposed while under ketamine/xylazine anesthesia using a closed-field speaker. Since anesthesia often reduces body temperature, which can reduce cochlear damage, mice exposed to 101 and 103 dB did not show permanent elevation of DPOAE thresholds (**Fig 1G,H**, cyan &magenta). However, mice exposed to 104 dB exhibited up to 30 dB of permanent threshold shift (orange). Once again, mean SR was unchanged relative to controls in all three groups (**Fig 2C,E**). A two-way ANOVA demonstrated no effect of exposure or frequency region (F_4,264_ = 0.8, p = 0.5; F_2,264_ = 0.3, p = 0.7). Thus, in our hands, the distribution of single-unit SRs was not obviously changed in the mouse IC following octave-band, 2-hr noise exposure regardless of anesthesia, intensity or laterality.

### Spontaneous rate was unchanged for all unit types: FRA classification

The central nucleus of the IC receives feed-forward axonal projections from nearly all brainstem auditory nuclei, and its physiological responses reflect this diversity of input (Ito et al., 2016). The presence and pattern of the inhibition in frequency response areas (FRAs) has been used to divide ICc neurons into response types V, I and O, which were suggested to arise due to dominant inputs from the medial superior olive, lateral superior olive, and dorsal cochlear nucleus, respectively (Ramachandran et al., 1999).

While there is anatomical evidence for division of inputs by source into “synaptic domains” at the circuit level (Ito et al., 2016), there is likely significant overlap between inputs from different sources at the regional level (Cant, 2005; Schofield, 2005). Synaptic integration of IC neurons by a heterogeneous mixture of inputs is further supported by an analysis of a large dataset of FRAs that indicates that response types form a continuous distribution rather than falling into discrete classes (Palmer et al., 2013).

Although the distribution may be continuous, it still may be informative to classify units, as such division has revealed class-specific changes to SR at more damaging exposure levels (Ropp et al., 2014). Therefore, we used FRAs to define three classes based on the bandwidth of excitation and the presence of on-CF inhibition (Egorova et al., 2001; Ma et al., 2006). “Broad” units had a V-shaped excitatory area and a wide bandwidth on both the low *and* high-frequency sides, with a total bandwidth at 10 dB above threshold (BW_10_) of at least one octave (**Fig S1A, Fig S2B**). “Narrow” units had an excitatory area ranging from V- to I-shaped with a particularly narrow bandwidth on the high-frequency side and a BW_10_ of less than one octave (**Fig S1B,C**). “Non-monotonic” units also had BW_10_ less than one octave and further demonstrated on-CF inhibition at high levels (**Fig S1D**). Inhibition was typically moderate (E/I Bal > 0), but in some cases was strong, causing the rate to CF tone to drop below SR (E/I Bal <0; **Fig S2A**; see Methods). The frequency extent of inhibition varied, causing upward-tilting, downward-tilting, and Type-O FRA shapes (Palmer et al., 2013). About 10% of units did not fit into any type and were classified as “Other”. These included units that were purely inhibitory, had multiple separate excitatory areas, or had CFs more than ½ octave removed from the local multi-unit CF.

Summed across all groups, the population was 55% narrow, 30% non-monotonic, 5% broad, and 10% other. Averaging within each anesthesia group and frequency band, and comparing across exposure condition (**Fig 3A-D, bottom**) revealed only small differences in the proportions of different classes, and a χ^2^ analysis did not reveal any significant differences due to exposure (Anesthetized [<16 kHz: 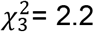, p=0.1, 16-32 kHz: 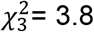, p=0.05, >32 kHz: 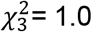, p=0.3]; Awake [<16 kHz: 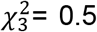, p=0.5, 16-32 kHz: 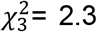, p=0.1, >32 kHz: 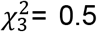, p=0.4]) or anesthesia (Control [<16 kHz: 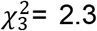, p=0.13, 16-32 kHz: 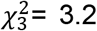, p=0.08, >32 kHz: 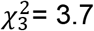, p=0.05]).

**Figure 3.**
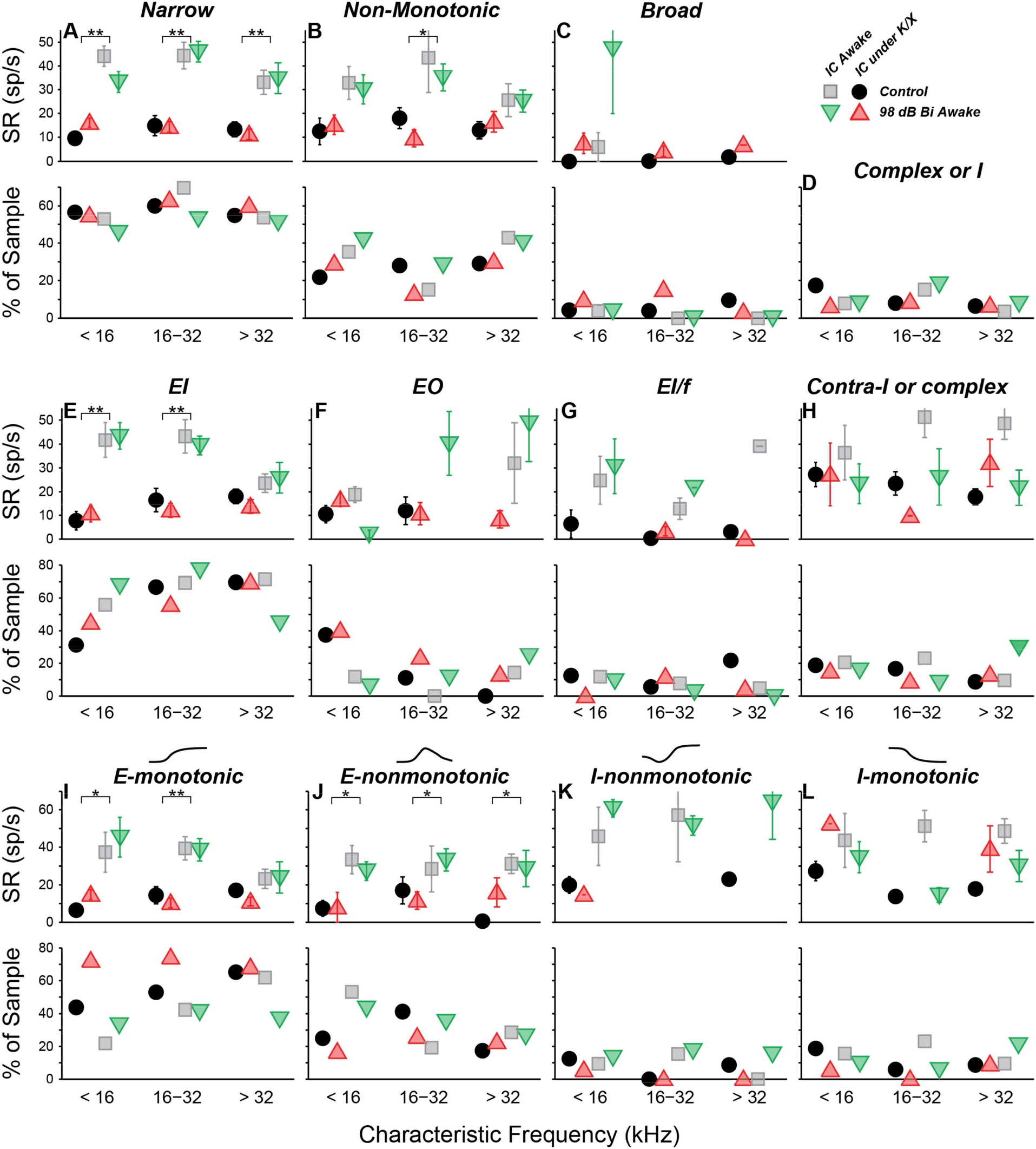
SRs in isolated IC neurons are unaffected by a neuropathic 98 dB noise exposure, even after segregating for response type. Three classification strategies are used: response area (**A-D**), E/I combination (**E-H**), or contralateral rate-level monotonicity (**E-I**). For each strategy, the top row shows mean SRs (±SEMs), and the bottom row shows percent of population. **A-C:** Units classified by response-area shapes - see Figures S1 and S2 for further details. **D:** Percent of sample for neurons unclassifable by response area. **E-H** Units classified by response to noise: **E:** Ipsi excitatory / contra inhibitory (El). **F:** Ipsi excitatory / contra no response (EO). G: Ipsi excitatory / contra inhibitory with binaural facilitation (El/f). **H:** Ipsi inhibitory / contra inhibitory (II), Inhibitory/No Response (IO), or complex unit types. **I-L:** Units classified by response to contralateral noise only, in order of increasing presence of inhibition: Excitatory-monotonic (**I**), Excitatory-nonmonotonic (**J**), Inhibitory-nonmonotonic (**K**), and Inhibitory-monotonic (**L**). Symbol types are as described in Figure 2.

A three-way ANOVA confirmed what is visually apparent: there were no within-type effects of exposure on SRs (**Fig 3A-C**). Among narrow units, the effect of anesthesia was highly significant (F_1,228_ = 107, p<<0.001), but effects of exposure and frequency bin were not (F_1,228_ = 0.1, p=0.8; F_2,228_ = 2.7, p=0.07). The same was true for non-monotonic units (F_1,117_ = 18, p<<0.001; F_1,117_ = 0.3, p=0.6; F_2,117_ = 0.8, p=0.5). The number of units in the broad and other categories was too small to make a meaningful comparison of SRs.

Narrow monotonic units are typically subdivided into Type V and Type I units on the basis of the bandwidth of excitatory tuning and presence of inhibitory sidebands (Ramachandran et al., 1999). In this dataset, 75% of narrow units had inhibitory sidebands (ranging from barely-detectable to substantial) and 10% had no detectable sidebands (**Fig S1B,C**). The remaining 15% had SRs near zero; assessment of sidebands in these neurons would require a two-tone stimulus paradigm, which was not collected. SR was not affected by exposure within any of these subgroups (data not shown).

### Spontaneous rate was unchanged for all unit types: binaural noise classification

Although IC units typically respond most robustly to contralateral sounds, their binaural response properties can be used to infer their dominant sub-collicular inputs. Canonically, neurons excited by both contralateral and ipsilateral sound (EE) receive dominant inputs from the medial superior olive, whereas neurons excited by contralateral sounds and inhibited by ipsilateral sound (EI) receive dominant inputs from the lateral superior olive, and neurons unaffected by ipsilateral sound (EO) receive dominant inputs from the cochlear nucleus (Davis et al., 1999). We constructed binaural-noise response areas by playing broadband noise to both ears simultaneously, and randomly varying the level in each ear. The majority of neurons (62%) were excited by contralateral noise and inhibited by ipsilateral noise (EI; **Fig S3A**). Neurons that were inhibited by contralateral noise at low levels, but excited at higher levels were also included in this group (e.g. **Fig S3H**). The strength of both ipsilateral and high-level contralateral inhibition varied substantially in this group. Consistent with the high-frequency hearing and underdeveloped medial superior olive in mouse (Irving and Harrison, 1967; Ollo and Schwartz, 1979), no EE neurons were encountered. While there were no neurons driven by ipsilateral noise alone, about 8% of neurons were *facilitated* by ipsilateral noise (EI/f; **Fig S3C,D**; Park and Pollak, 1993; Davis et al., 1999), and another 15% were driven by contralateral noise and unaffected by ipsilateral noise (EO; **Fig S3E&F**). The remaining 15% were inhibited by contralateral noise (**Fig S3G**), were not driven by noise, or had more complex responses. We calculated the proportions in each class within each anesthesia group and frequency band, and compared them across exposure condition (**Fig 3E-H**). A *χ*^2^ analysis for differences in the population statistics was significant in the >32 kHz anesthetized, 16-32 kHz awake, and >32 kHz awake cases (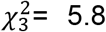, p=0.015; 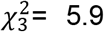, p=0.015; 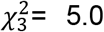, p=0.026 respectively). However, samples sizes were small, and after applying Yate’s correction for small samples no significant differences remained (p=0.09; p=0.09; p=0.13).

There were no within-class effects of exposure on SRs (**Fig 3E-H**). Among EI units, the effect of anesthesia was highly significant (F_1,174_ = 47, p<<0.001), but effects of exposure and frequency range were not (F_1,174_ = 0.1, p=0.8; F_2,174_ = 2.4, p=0.09). The number of units in the other categories was too small to make a meaningful comparison of SRs.

Because the number of neurons in the EI group was so large, this class was further divided based on the shape of the rate-level functions to contralateral noise: excitatory-monotonic neurons had strictly-increasing rate with increasing level, excitatory-nonmonotonic neurons had functions that rose and then fell at high levels, inhibitory-nonmonotonic neurons had functions that fell and then rose at high levels, and inhibitory-monotonic neurons had functions that strictly decreased. No significant exposure-related differences were found in either the proportion of neurons, or the SR, within each class.

### The slope of tone-evoked rate-vs-level functions was elevated in neuropathic mice

Noise exposure can enhance sound-evoked responses in the ascending auditory pathways. Prior studies have reported increased sound-evoked responses following noise exposure in the ventral cochlear nucleus (Cai et al., 2009), IC (Salvi et al., 1990), and auditory cortex (Resnik and Polley, 2017; reviewed in Auerbach et al., 2014). . Increased intensity of sound-evoked EEG and fMRI signals have been reported in humans with hyperacusis, even after controlling for audiometric thresholds (Gu et al., 2010, 2012). We therefore searched for neuronal hyperactivity by quantifying responses to CF tones obtained in the measurement of FRAs. Rate-level functions were fit with the four-parameter model of Sachs and Abbas (1974), or a seven-parameter sum of two such models (see Methods & **Fig S1**); the excitatory portion of this fit was then used to calculate threshold, maximum driven rate, and slope (**Fig 4**). In most comparisons there was no effect of exposure. However, in the region of maximal neuropathy (16-32 kHz), slopes were elevated by 65% in awake mice (**Fig 4I**, p=0.037, rank-sum test). Strikingly, slopes were not elevated under anesthesia within this (or any) frequency range, suggesting that hyperactivity may be modulated by anesthesia.

**Figure 4.**
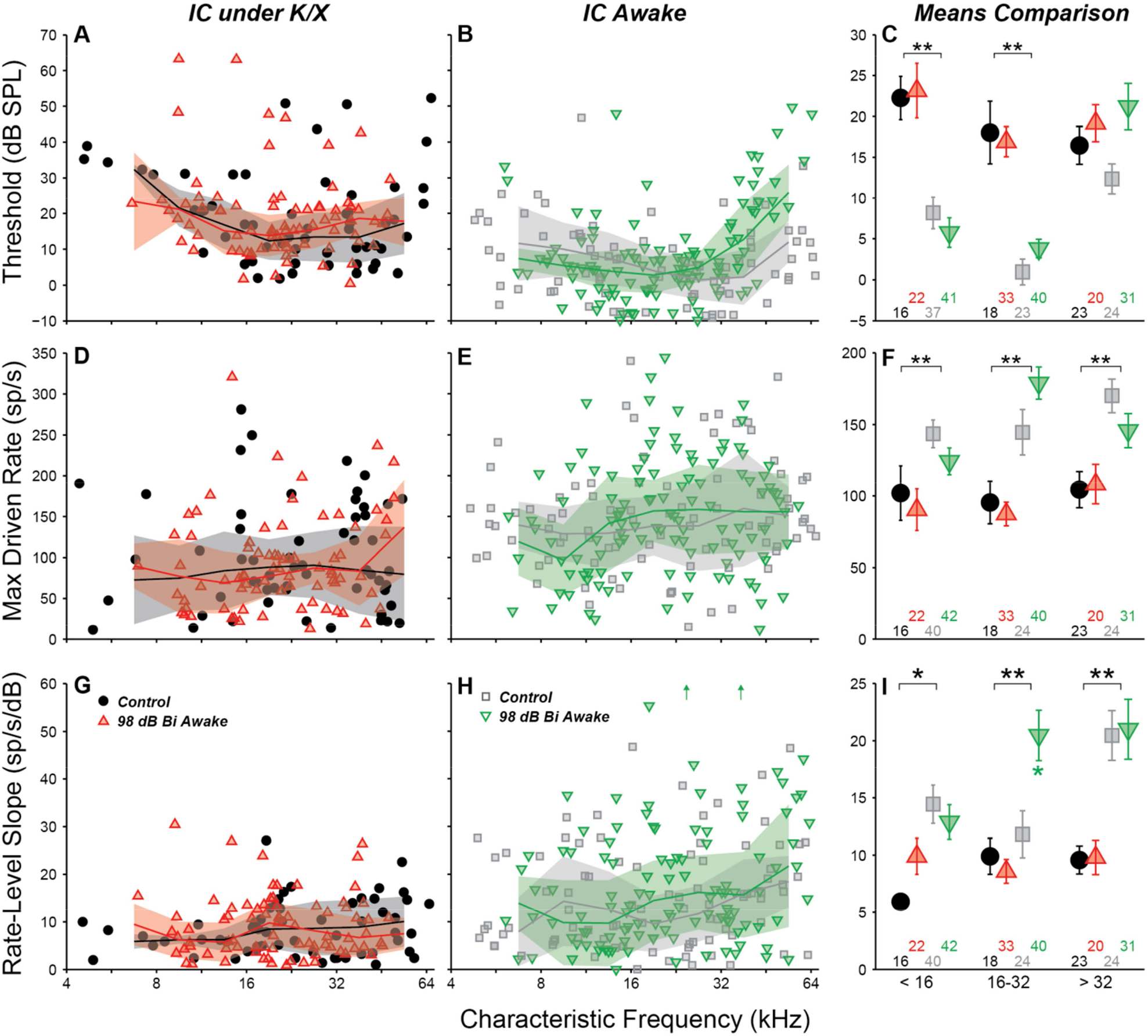
Responses to CF tones in isolated IC neurons from anesthetized (**A,D,G**) or awake (**B,E,F**) animals. The three rows show different metrics: threshold (**A,B,C**), maximum driven rate (maximum rate minus SR; **D,E,F**) and the slope of the rate-level function (**G,H,I**). **C,F,I:** Mean value (± SEM) for each parameter, binned into three CF regions as indicated. Units with inhibitory or complex response areas are not included. Off-axis values in (up arrows in **H**) were 64.2 and 60.9 from left to right. Lines and shading indicate median and interquartile range, calculated over a 1 octave sliding window. Black asterisks indicate significant effects of anesthesia; green asterisks (panel I only) indicate significant effects of noise exposure: p<0.05 (single), p<0.001 (double) by a rank-sum test with Bonferroni-Holm correction.

Anesthesia had a strong effect on all three parameters. Thresholds were significantly elevated by anesthesia in the non-neuropathic and neuropathic regions (p<<0.001, p<<0.001, rank-sum test). As observed with SRs, maximum driven rates were reduced by anesthesia in all frequency regions (p<0.001, p<<0.001, p<0.001). Rate-level function slopes were also significantly reduced across all CFs (p=0.006, p=0.001, p<<0.001).

Separating into unit classes revealed that slope elevations in the composite means were driven by increased slopes in the rising portion of non-monotonic neurons rate-level functions (**Fig 5A-C**). Exposure did not have a significant effect on threshold or maximum driven rate for any response type (data not shown). However, mean rate-level functions demonstrated a trend towards elevated maximum driven rates for both narrow and non-monotonic neurons in the neuropathic region (**Fig 5D,E**). This trend was not present in the other frequency regions (data not shown).

**Figure 5.**
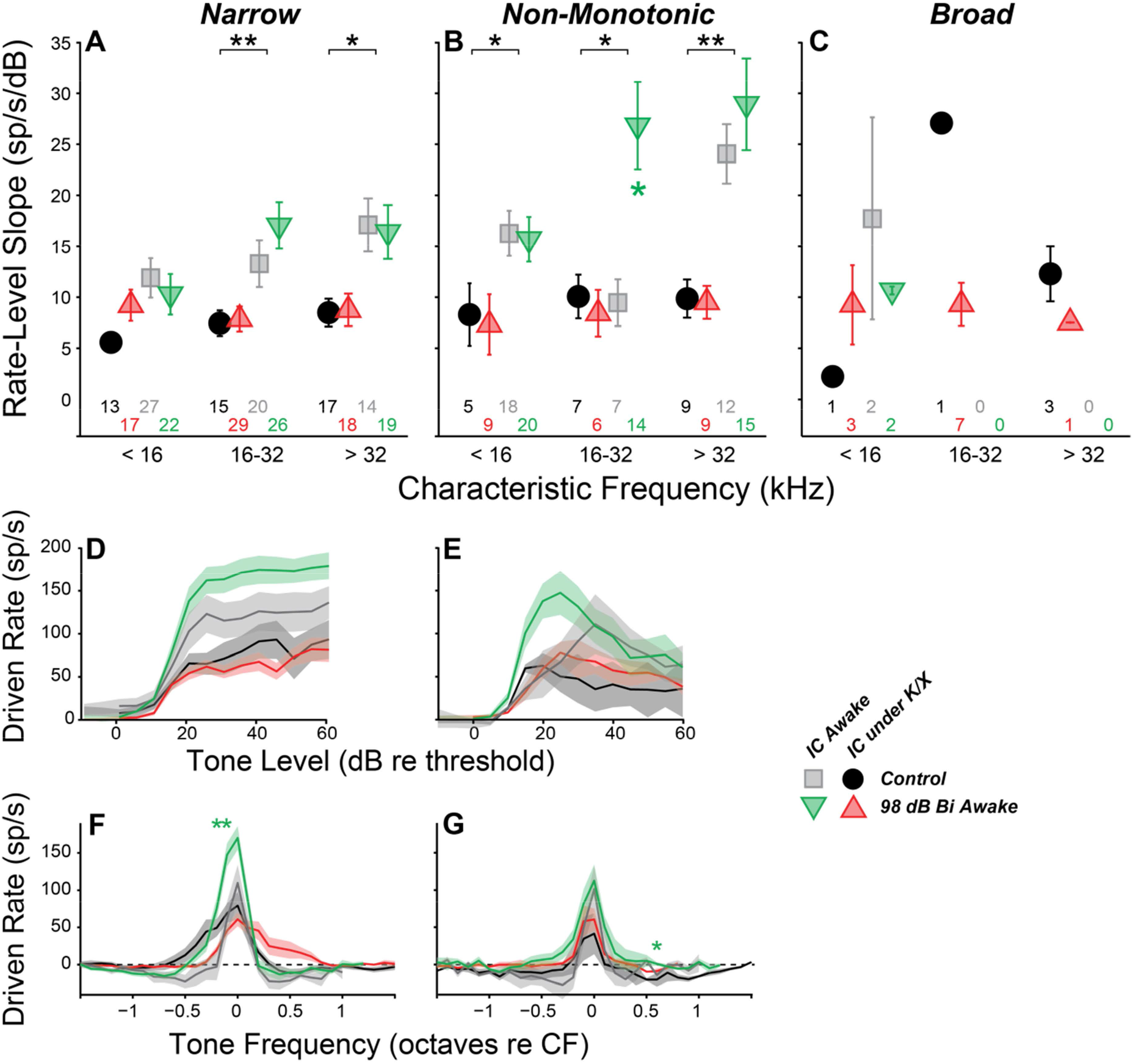
Slope increases in rate-level functions to tones are seen only in non-monotonic IC neurons from neuropathic regions (16-32 kHz). **A,B,C:** Mean slope (± SEM) of rate-level functions to tones at CF in single units classified by their frequency response areas as broad (**A**), narrow (**B**), or non-monotonic (**C**). **D-G:** Mean tone-driven response for narrow (**D,F**) and non-monotonic (**E,G**) neurons with CFs between 16 and 32 kHz: either rate-level functions to CF tones (D,E) or rate-frequency sweeps averaged at levels 20, 25, and 30 dB above threshold. Black asterisks indicate significant effects of anesthesia, as described in Figure 4; green asterisks indicate significant effects of exposure on IC responses (p<0.05 (single), p<0.001 (double) by Tukey-Kramer-corrected post-hoc test following a 3-way ANOVA). There were no significant effects of exposure on IC responses in anesthetized animals.

### Inhibitory side bands were reduced in neuropathic mice

Responses to off-CF tones were quantified by plotting driven rate vs. frequency for tones 20 - 30 dB above threshold. In the neuropathic region of awake mice, driven rate was elevated at frequencies above and below CF. This was apparent when pooling all neurons (data not shown), but most prominent in neurons with a narrow tone-response type (**Fig 5F vs. G**). Notably, while mean driven rate dropped below 0 in control neurons, presumably reflecting sideband inhibition, driven rate remained positive in exposed neurons (**Fig 5F,G**). This reduction in off-CF inhibition could be related to the elevated rate-level slopes at CF. Significant effects of exposure were not observed at other frequency regions (data not shown).

### Responses to noise were unchanged in neuropathic mice

Rate-level functions to contralateral noise were also extracted from the binaural noise response maps (**Fig S3**). In the ensemble averages, there were no significant effects of exposure on threshold, maximum driven rate, or slope (**Fig 6**). No effects of exposure were revealed by classifying neurons by tone-, binaural-noise, or contralateral-noise response types (data not shown).

**Figure 6.**
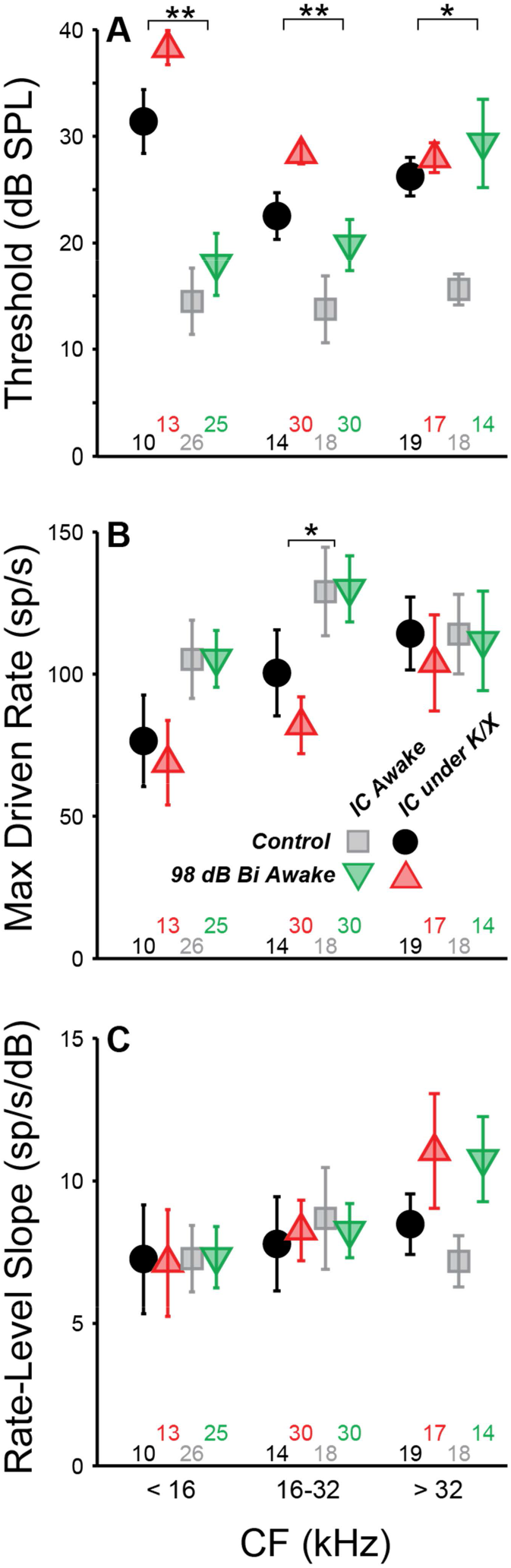
IC Single-unit responses to noise were unchanged following neuropathic damage. Mean threshold (**A**), maximum driven rate (**B**), and slope of the rate-level functions (C) for contralateral broadband noise. Black asterisks indicate significant effects of anesthesia as described in Figure 4. There were no significant effects of exposure.

### Exposure effects on multi-unit responses were different than those seen in single-units

Single-unit responses in the IC are inherently variable, given the complex mixture of response types and varied input circuits. This heterogeneity in normal response profiles could hide important effects of noise exposure on stimulus coding in the IC. As an alternative to single-unit approaches, some prior studies (Hesse et al., 2016) have relied on multi-unit activity (MUA), which can be easier to measure, and, by its nature, averages response of many adjacent neurons, especially at high stimulus levels. In the IC, multiunit FRAs are usually Type V, i.e. with minimal inhibitory sidebands and monotonic rate-level functions at CF. While fine detail is obscured in MUA measures, they may be useful in evaluating gross changes in IC responses due to noise exposure.

Here, MUA was quantified in two ways: 1) by counting spikes after thresholding the electrode signal at 3.5 standard deviations of the background noise (tMUA, **Fig 7A**) and 2) by measuring the envelope of the MUA (eMUA), without any thresholding step (**Fig 7B**) (Supèr and Roelfsema, 2005; Kayser et al., 2007; Choi et al., 2010). Following neuropathic exposure, spontaneous activity was slightly (but significantly) reduced in the neuropathic region by both measures in anesthetized mice. However, in awake mice, spontaneous activity in the neuropathic region was elevated 6% following exposure in the tMUA, but reduced 10% in the eMUA. We have no clear explanation for these divergent effects, but note they are much smaller than the elevations in MUA-based spontaneous activity following exposure reported by Hesse et al. 2016, which ranged from 130 to 600%, depending on frequency region.

**Figure 7.**
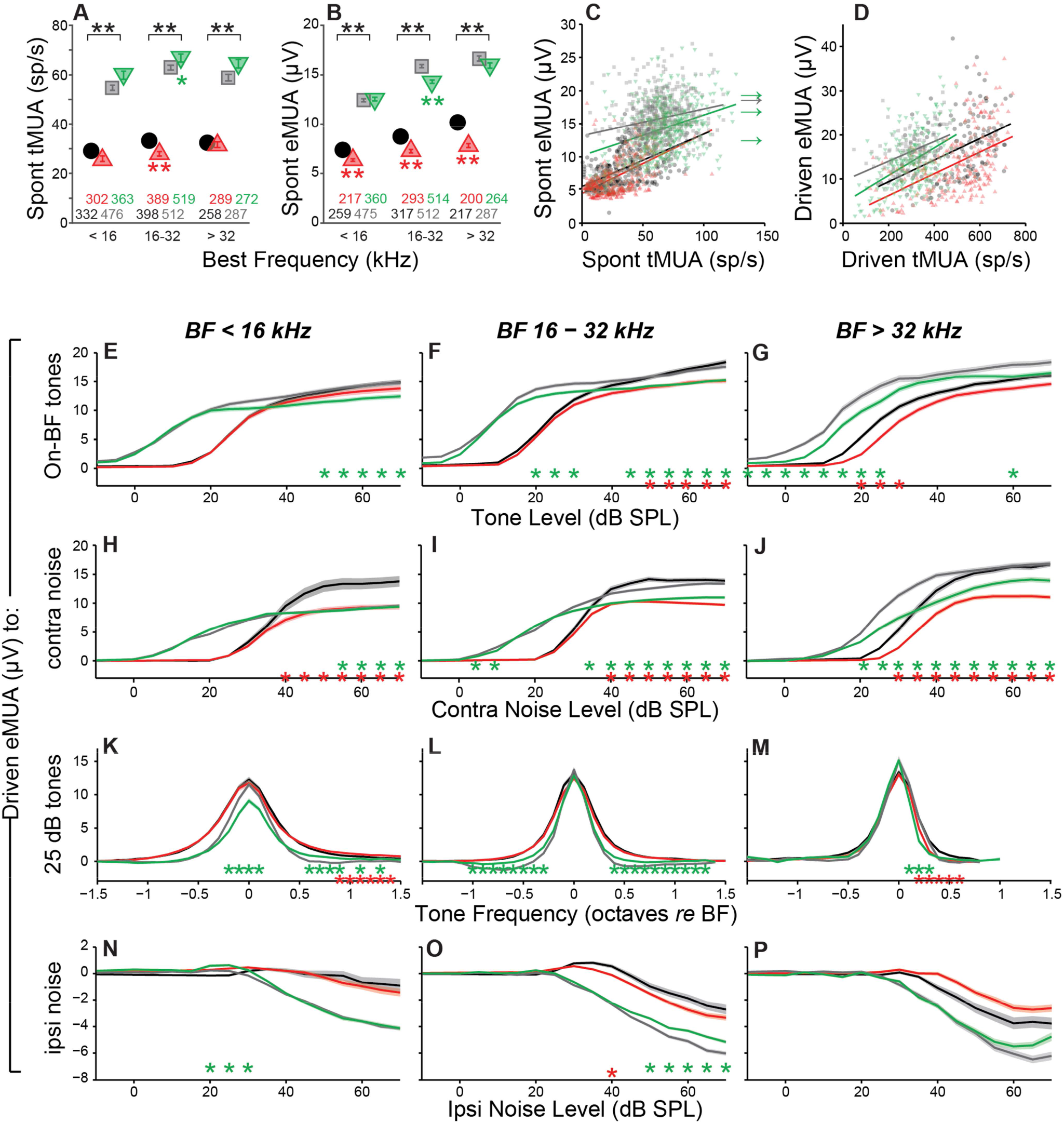
Multi-unit IC results do not mirror single-unit IC results. **A-B:** Mean spontaneous activity of multi-units for three BF bins (as indicated), as estimated by eMUA (**A**) or by tMUA (**B**). Asterisks indicate significant effects of anesthesia or noise exposure, as described in Figure 4. **C:** Multi-unit estimates of spontaneous activity compared for the two metrics for each site. **D:** Multi-unit estimates of driven rate for a BF tone 40 dB above threshold. **E-P:** Average multi-unit driven-rate vs. level or frequency for eMUA, as averaged over three CF regions (three columns) and for four different stimuli (four rows), as indicated. Stimuli were: BF tones (**E,F,G**); contralateral broadband noise (**H,I,J**), tones at 20, 25, and 30 dB above threshold (**K,L,M**) or ipsilateral noise with concurrent 30 dB SPL contralateral noise (**N,O,P**). Asterisks indicate significant effects of anesthesia (black) or noise exposure (green and red) as described in Figure 5.

We next compared the two MUA metrics at each recording site (**Fig 7C,D**). The two measures were moderately correlated for tone-driven activity, regardless of anesthesia (r^2^ = [0.19, 0.25, 0.17, 0.22] for [control k/x, exposed k/x, control awake, exposed awake] respectively, all p<<0.001), but poorly correlated for spontaneous activity in awake mice (r^2^ = [0.30, 0.29, 0.05, 0.11] for [control k/x, exposed k/x, control awake, exposed awake] respectively, all p<<0.001). In a pattern opposite to that in single units, mean tone-evoked tMUA was 35% lower in awake mice. This decrease may not reflect underlying single-unit hypoactivity, but a difference in the (3.5 SD) threshold for spike detection: median threshold was 70% higher for awake recordings than for anesthetized recordings (data not shown), presumably because the SD is computed from the electrode signal during silence, and single-unit SRs were elevated in awake mice *re* anesthetized animals (**Fig 2D**).

Since quantification of eMUA does not require thresholding, we evaluated the effect of neuropathic exposures on tone- and noise-driven eMUA. Responses to tones at the best frequency (BF) were reduced at moderate-to-high levels in the non-neuropathic region of awake mice, and in the neuropathic region of both awake and anesthetized mice (**Fig 7E-G**). This trend was visible in single units, but more robust and highly significant here. Unlike in single units, noise-driven eMUA was reduced at moderate-to-high levels in the non-neuropathic region of anesthetized mice, and in the neuropathic region of both awake and anesthetized mice (**Fig 7H-J**). These reductions might be caused by a direct effect of cochlear neuropathy, the loss of high-threshold, low-SR auditory-nerve fibers.

Driven responses to tones with frequencies both above and below BF were elevated in the neuropathic region of noise exposed mice (**Fig 7K**). Like the on-BF changes, this effect was also visible in single units, but was more robust in the eMUA. A potentially related loss of inhibition was observed as a reduction in the suppression of eMUA by ipsilateral noise (**Fig 7N-P**). This trend was visible but not significant in single-unit responses (data not shown).

## DISCUSSION

Noise exposure is the most common cause of chronic tinnitus and hyperacusis in human subjects, and these perceptual anomalies often begin immediately post exposure (Meikle, 1997; Holgers and Pettersson, 2005; Daniel, 2007). They can last indefinitely and can be elicited whether or not there is a noise-induced PTS (Axelsson and Ringdahl, 1989; Shargorodsky et al., 2010). The tinnitus percept is often tonal in nature, and, when there is PTS, the pitch often falls within the PTS region (Sereda et al., 2011, 2015; Schecklmann et al., 2012), consistent with models suggesting it arises due to compensatory homeostatic plasticity in response to reduced peripheral input (Schaette and Kempter, 2012; Noreña and Farley, 2013; Auerbach et al., 2014).

Since tinnitus is the percept of sound in the absence of sound, many neurophysiological studies have looked for noise-induced changes in spontaneous activity in auditory neurons from cochlea to cortex. Although some have suggested that changes to discharge synchrony across neuronal populations or increased bursting activity in the SR might be key to tinnitus generation (reviewed by Roberts et al., 2010), most studies have concentrated on SRs.

In the auditory nerve, SR is either unchanged or reduced by acoustic overexposure (Liberman and Kiang, 1978; Liberman and Dodds, 1984; Furman, 2013). Auditory-nerve SR is created at the inner hair cell synapse (Rutherford and Moser, 2016), thus, loss of inner hair cells will silence all the fibers that formerly contacted them (Liberman and Kiang, 1978). Correspondingly, loss of outer hair cells does not directly affect SR (Dallos and Harris, 1978; Schmiedt et al., 1980), although it will cause PTS due to loss of the mechanical amplification that outer hair cells provide (Ryan and Dallos, 1975). Damage to, or loss of, inner hair cell stereocilia, a common result of acoustic overstimulation, also causes PTS, but also reduces auditory-nerve SR by inactivating mechanoelectric transduction channels, thereby reducing leak current, hyperpolarizing the hair cell, and reducing vesicle release at the synapse (Liberman and Dodds, 1984). The auditory-nerve synapses with inner hair cells are the most vulnerable elements to noise damage (Kujawa and Liberman, 2009), and synaptopathy silences those fibers that become disconnected from their peripheral targets. Because synaptopathy can be widespread in ears without hair cell loss or PTS, it might underlie the generation of noise-induced tinnitus in ears with a normal audiogram. Our major aim in the present study was to evaluate this notion.

Hyperacusis is the sensation that moderate-level sounds are intolerably loud, and the search for neurophysiological correlates have typically examined on rate-level functions to CF tones. At the auditory nerve level, as for SRs, noise damage generally reduces maximum discharge rates and the slopes of the tone-evoked rate-level functions (Liberman and Kiang, 1978; Heinz and Young, 2004).

Although noise damage invariably reduces SR and tone-evoked rates in the auditory periphery, it often elevates SR and tone-evoked rates in higher auditory centers (Eggermont and Roberts, 2004), presumably via re-adjustment of “central gain” (Schaette and Kempter, 2012; Noreña and Farley, 2013; Auerbach et al., 2014). Hyperactivity has been reported in the IC of several species following acoustic overexposures that were either tonal or broadband, unilateral or bilateral, and in awake or anesthetized animals (**Table 1**). SR elevations can be as large as 50 fold, can be observed as soon as a few hrs post-exposure, and can persist for at least a year (**Table 1**). While exposures used in most IC studies elicited > 30 dB of PTS, elevated SRs have been reported in cases with minimal PTS in both mice (Longenecker and Galazyuk, 2011; Hesse et al., 2016) and chinchillas (Bauer et al., 2008). However, the nature and degree of cochlear damage has rarely been reported, and the role of synaptopathy as a key elicitor has only recently been evaluated (Hesse et al., 2016).

**Table 1.**
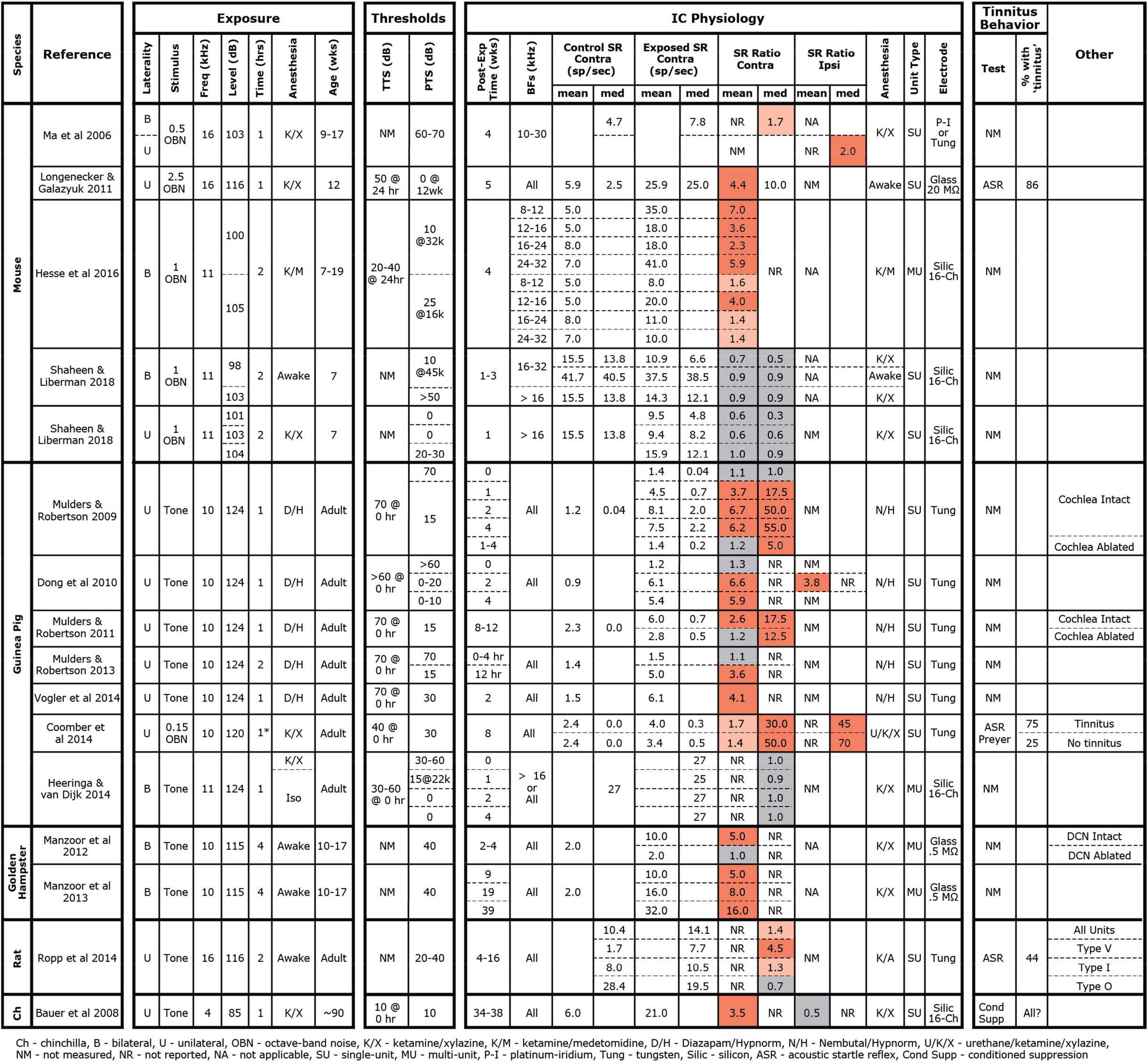
Summary of studies measuring SRs in the IC following noise exposure. Cells are colorized according to degree of noise-induced changes in SR.

### Effects of noise exposure on spontaneous rate

Here, bilateral exposure to octave-band noise at 98 dB SPL did not change population measures of SR in the IC, despite substantial permanent synaptic loss throughout a large portion of the basal cochlea (**Figs 1&2**). This result was replicated in recordings from both anesthetized and awake animals. Subdividing the neural population into response types did not uncover any class-specific SR changes. Thus, noise-induced synaptic loss does not appear to be sufficient to cause increased SRs in the mouse IC, at least in mice exposed at 7 wks of age.

We increased the noise level to produce a 50 dB PTS, thereby adding significant hair cell damage to the spectrum of cochlear pathology, but SRs remained unchanged (**Figs 1&2**, blue diamonds). Particularly large SR increases have been reported in IC following unilateral exposures: 10 fold in mice (Longenecker and Galazyuk, 2011) and 17-50 fold in guinea pigs (Mulders and Robertson, 2009, 2011). Therefore, we exposed 3 groups of mice unilaterally to 8-16 kHz noise at three different intensities, however SRs were not elevated in these groups either, regardless of the amount of PTS.

In interpreting results across studies, electrode style and reliance on single unit vs. multi-unit activity must be considered. Here, we got contradictory results when assessing data from the same penetrations as single-vs. multi-unit activity. Although single-unit records are likely more reliable, referring to **Table 1** shows that this distinction does not resolve the discrepancy. Similarly, some studies used micropipettes with fine tips (3 μm), while we used silicon electrodes with large contact areas (177 μm^2^). The smaller micropipettes may detect signals from smaller neural elements, including fibers of passage; thus different neural populations may be sampled. However, this distinction also does not resolve the discrepancy.

The presence of anesthesia during exposure can change cochlear vulnerability (**Fig 1**), and might also change exposure effects on central pathways. However, SR increases have been observed in animals exposed either awake or anesthetized (see **Table 1**), whereas we failed to see SR increases in animals exposed either way. In anesthetized mouse IC, Hesse et al. 2016 measured multi-unit SRs and reported a 2-7 fold increase 4 wks following an 8-16 kHz, bilateral, 100 dB exposure of anesthetized animals. Computing multi-unit SRs in a similar way, we observed slight decreases at 1-3 wks post-exposure (**Fig 7**).

Post-exposure survival could be an important variable, as noise-induced SR changes are likely dynamic in the hours and days after damage. In our study, SRs were measured at 1-3 wks post-exposure, a shorter time period than the other mouse studies (**Table 1**). In a recent study in awake mice of exactly the same noise exposure used here, SRs in primary auditory cortex were unchanged 1 day post-exposure, but elevated 3-fold by 4 days and back to baseline by 1 wk, where they remained for next 6 wks post exposure (Resnik and Polley, 2017). The lack of SR changes observed here in the IC at 1-3 wks post-exposure are consistent with that pattern. Noise-induced changes in SR appear stable at longer post-exposure times. In noise-exposed guinea pigs, SRs in the IC are unchanged immediately post-exposure, but elevated by 12 hrs, where they remain elevated for at least 8 wks post-exposure (Mulders and Robertson, 2009, 2011, 2013). The sources of elevated SRs may migrate centrally at longer post-exposure times: in guinea pigs, cochlear ablation < 4 wks post-exposure returns SRs in the IC to control levels, whereas ablation > 8 wks post-exposure does not (Mulders and Robertson, 2009, 2011). However, there are no studies of SR post-exposure dynamics suggesting that elevated SRs would have developed in the present study if survival had been extended beyond 3 wks.

Age at exposure could also be an important variable, as cochlear vulnerability to noise decreases dramatically in mice between 8 and 16 wks (Kujawa and Liberman, 2006), and adolescent central auditory pathways may react differently to noise than in the adult. In our study, mice were exposed at 7 wks, an age we have used for many recent studies of cochlear synaptopathy, because it is after the onset of sexual maturity (~ 4-5 wks) but before the period where vulnerability is changing rapidly with age (thereby introducing a potential source of variability). As seen in Table 1, this is a younger age than that in the other mouse studies, and, although cross-species age comparison is difficult, may also represent a relatively younger age than in all other studies summarized there. Changes in excitatory/inhibitory balance that must underlie noise-induced hyperactivity may be more effectively rebalanced in the young brain than in the adult. It is interesting, in this regard, that tinnitus is much less common in younger people than in adults (Martinez et al., 2015).

Another intriguing clue to the discrepancy between our results and prior studies is that our control SRs were significantly higher than in other mouse studies, e.g. 2.5X higher than others in anesthetized mice (Ma et al 2006; Hesse et al 2016) and 6X higher than others in awake mice (Longenecker and Galazyuk, 2011). This, in turn, suggests a difference in pre-exposure excitatory/inhibitory balance in the IC, which is interesting given reports that the emergence of noise-induced SR elevations and tinnitus can also depend on pre-exposure state. Given identical noise exposure, gerbils with low pre-exposure spontaneous and sound-evoked neural activity in the auditory cortex and brainstem develop “tinnitus” and cortical SR elevations, while those with higher pre-exposure activity do not (Ahlf et al., 2012; Tziridis et al., 2015). The cause of the differences in pre-exposure state between the two groups is unclear, but may be critically important in understanding the genesis of tinnitus.

### Effects of noise exposure on sound-evoked responses

Here, we saw no significant noise-induced effects on the maximum tone-evoked rates of IC neurons, however the slopes of rate-level functions to CF tones were elevated in a subset of neurons (**Figs 4 & 5**). This effect was seen only in neurons with CFs in the affected region (16-32 kHz), strongly suggesting it might be caused by the cochlear synaptopathy *per se*. The effect was also seen only in units with non-monotonic rate-level functions, suggesting that only a subset of the central circuitry appears to compensate for a reduced peripheral drive by increasing its overall gain. In the face of reduced peripheral input, the balance of excitation and inhibition may be adjusted to maintain detectability. Since non-monotonic neurons receive substantial on-CF inhibition, the shape of their rate-level functions may be particularly sensitive to this balance. Changes to the excitatory/inhibitory balance were also observed in response to off-CF tones (**Fig 5**) and to binaural noise (**Fig 6**). In the chinchilla IC, elevated evoked rates were observed in a larger proportion of non-monotonic neurons than monotonic neurons immediately following exposure to a high-intensity, above-CF tone (Wang et al., 1996). The authors suggest that this difference could be due to a reduction of lateral inhibition from the exposed region. Ketamine, an NDMA receptor agonist, reduces excitatory drive, which could mask changes to the excitatory/inhibitory balance and may be the reason that slopes, in our study, were unaffected by noise exposure in anesthetized mice (**Fig 5**).

In mice exposed to the same traumatic noise used here, both SR and tone rate-level slopes were transiently increased in auditory cortical neurons 3-4 days post exposure, recovering to baseline by 1 wk (Resnik and Polley, 2017). Again using the same noise exposure, calcium imaging of cortico-collicular neurons revealed increased rate-level slopes 2 days post exposure that remained elevated to at least 2 wks (Asokan et al., 2018). Here, we didn’t track neuronal responsiveness at post-exposure times < 1 wk, when larger effects might have been seen. More extensive and long-lasting changes to the excitatory/inhibitory balance were observed following near-complete cochlear synaptopathy by cochlear administration of a Na/K pump blocker (ouabain): despite a massive loss of peripheral input, tone rate-level slopes recovered in the IC and were elevated in the auditory cortex by 30 days post-treatment (Chambers et al., 2016). Clearly, the nature and extent of central hyperactivity following peripheral lesion is a complex function of the lesion severity, the region of the auditory pathway under study, the postexposure time and the baseline state at the time of exposure (**Table 1**).

### Tinnitus, hyperacusis, and central gain

A number of tests are used to infer the presence of tinnitus in animals. One is based on prepulse inhibition (PPI) of the acoustic startle reflex, where the “prepulse” is a silent gap in an otherwise continuous background noise. The idea is that a tinnitus percept makes the noise gap less salient and reduces gap-induced PPI of startle (Turner et al., 2006). In a prior study of mice, evaluated 1-10 wks after exposure to the same traumatic noise used here, animals showed reduction of gap-induced PPI when the background was narrow-band noise centered in the synaptopathic region (Hickox and Liberman, 2014). This might suggest a tonal tinnitus near 32 kHz. However the effect was seen only with no delay between gap offset and startle onset, not with an 80 msec gap-startle delay. This dependence on stimulus timing casts doubt as to whether the tinnitus was really “filling the gap” and thus on the relevance of the gap-startle test to tinnitus. It also leaves open the question of whether this synaptopathic exposure produces tinnitus in mice.

Studies of SR in the dorsal cochlear nucleus and thalamus report elevations only in animals with reduced gap-induced PPI (Dehmel et al., 2012a; Li et al., 2013; Kalappa et al., 2014), while SRs in the IC and auditory cortex are elevated regardless of the results of a gap-induced PPI assay (Engineer et al., 2011; Coomber et al., 2014; Ropp et al., 2014). Elevated SRs have been reported in the dorsal cochlear nucleus following synaptopathic noise exposures that cause no PTS in chinchillas (Bauer et al., 2008) and guinea pigs (Koehler and Shore, 2013). However, based on either gap-indiced PPI (Koehler and Shore, 2013) or an operant conditioning test of tone perception (Bauer et al., 2008), only some exposed animals showed tinnitus. Together, these results suggest that synaptopathy alone is not sufficient to cause either elevated SRs or tinnitus. However, since all the studies in Table 1 likely cause significant synaptopathy, it remains possible that synaptopathy is a key factor in the generation of noise-induced tinnitus and SR elevation. Whether the reductions in ascending auditory drive from the periphery are translated into chronic hyperactivity in higher auditory centers likely depends on the intersubject differences in circuit plasticity within the central circuits themselves.

In a prior study, mice exposed to the same noise used here, showed significant hypersensitivity to moderate-level sound, as seen in decreased thresholds and increased amplitudes of the acoustic startle reflex, as measured without PPI (Hickox and Liberman, 2014). Since the IC is a dominant contributor to these responses (Fendt et al., 2001), increased IC rate-level slopes may be key to the increased startle responses associated with cochlear synaptopathy, which in turn may be related to the decreased noise-level tolerance in humans with acoustic overexposure.

---

**Figure S1.**
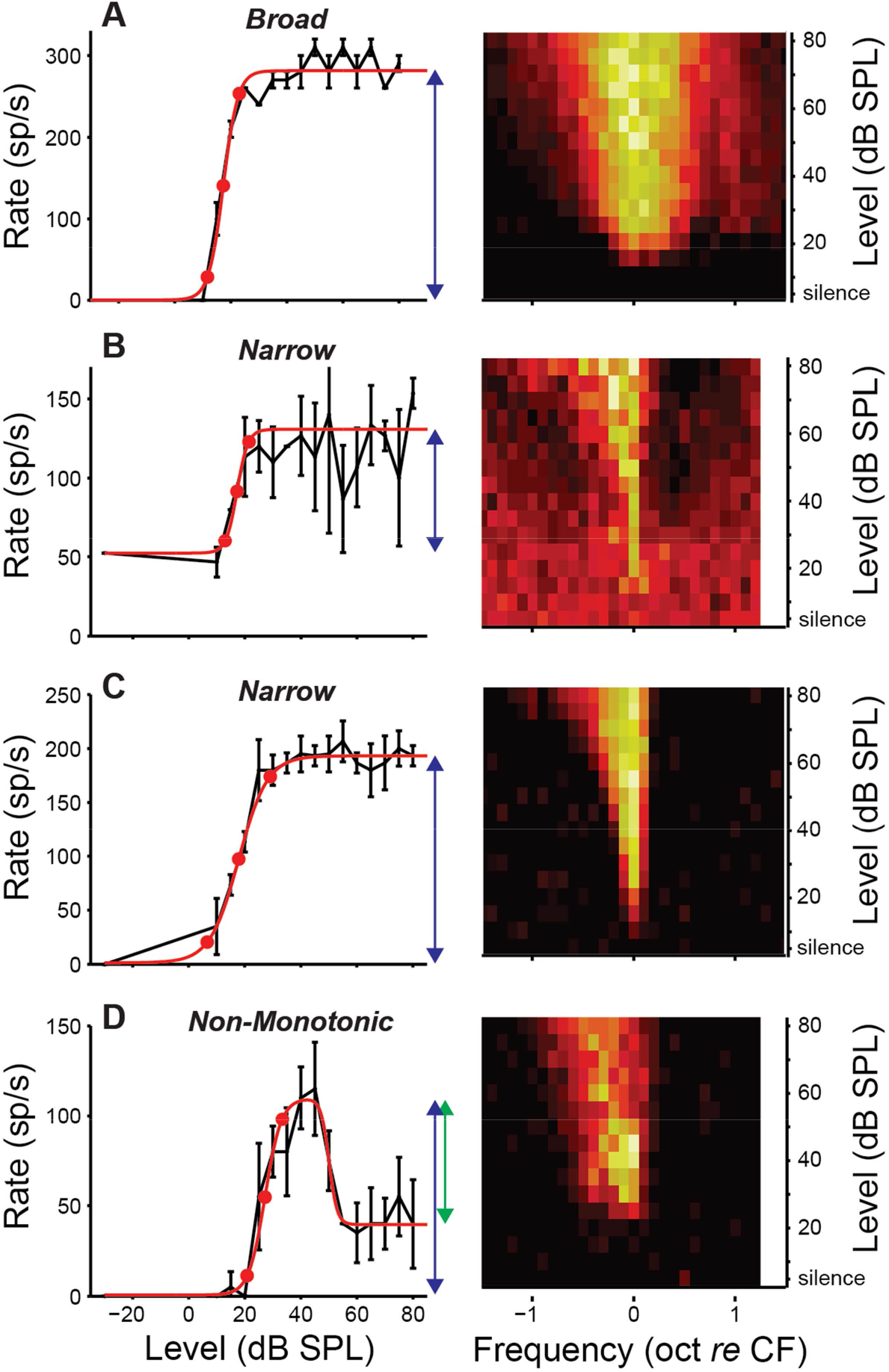
Unit types based on frequency response areas in isolated IC neurons. **A-D, Right**, Frequency response areas for four examplars from controls. Color indicates firing rate, normalized to maximum rate. **A-D, Left**, Rate responses to a tone at CF. Black points indicate mean rate and standard error. Red line indicates best-fit function with 10, 50, and 90% of the excitatory driven rate range (E_D_, blue arrow) indicated by red circles. Inhibitory driven rate (I_D_) is indicated in **D** by green arrow. E/I Balance in this unit: (E_D_ - I_D_) / (E_D_ + I_D_) = 0.36.

**Figure S2.**
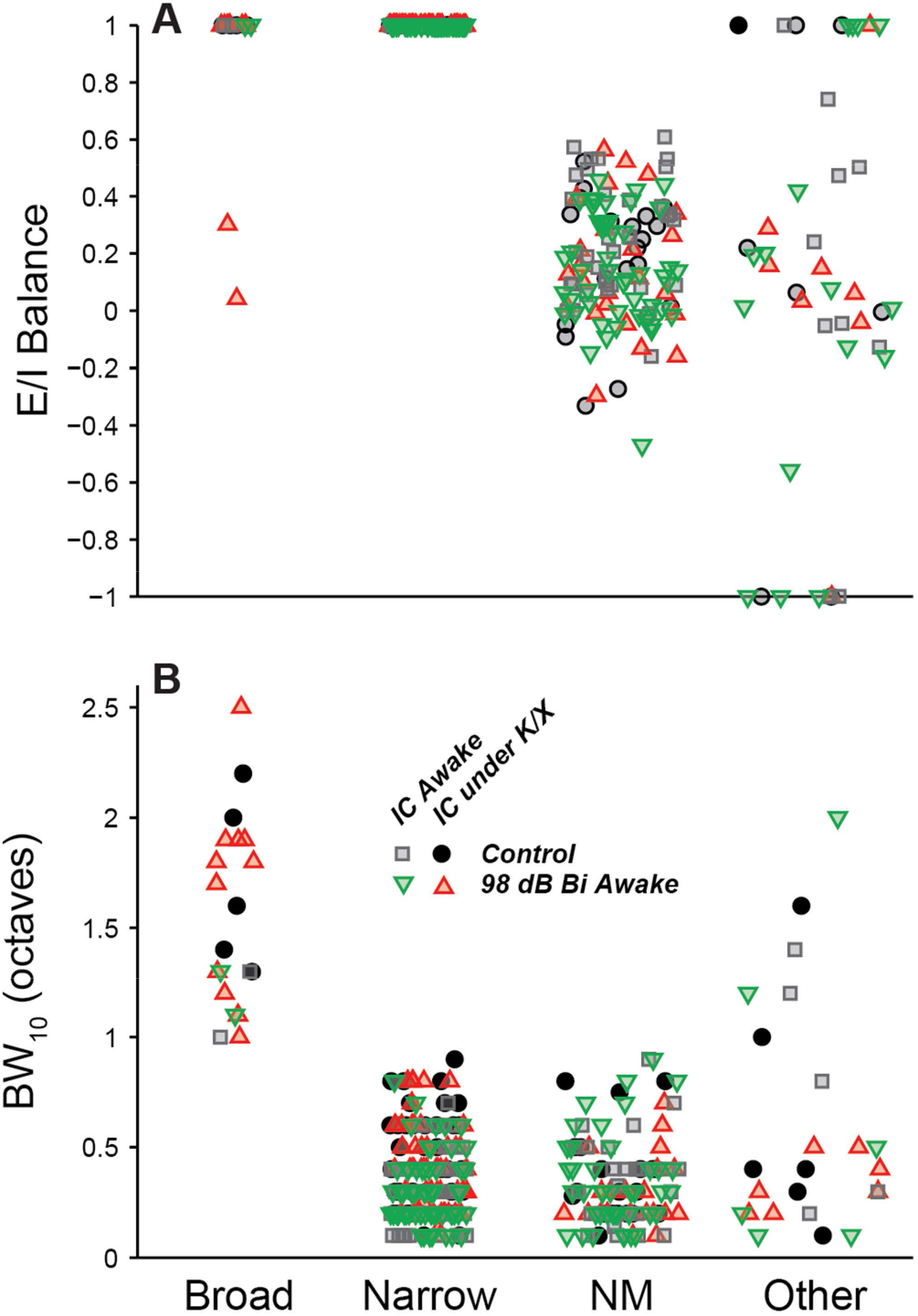
Metrics used for unit typing by frequency response area. **A:** Excitatory/Inhibitory balance (see Figure S1), the difference between excitatory and inhibitory drive normalized by their sum. **B:** The bandwidth of the excitatory response at 10 dB above threshold.

**Figure S3.**
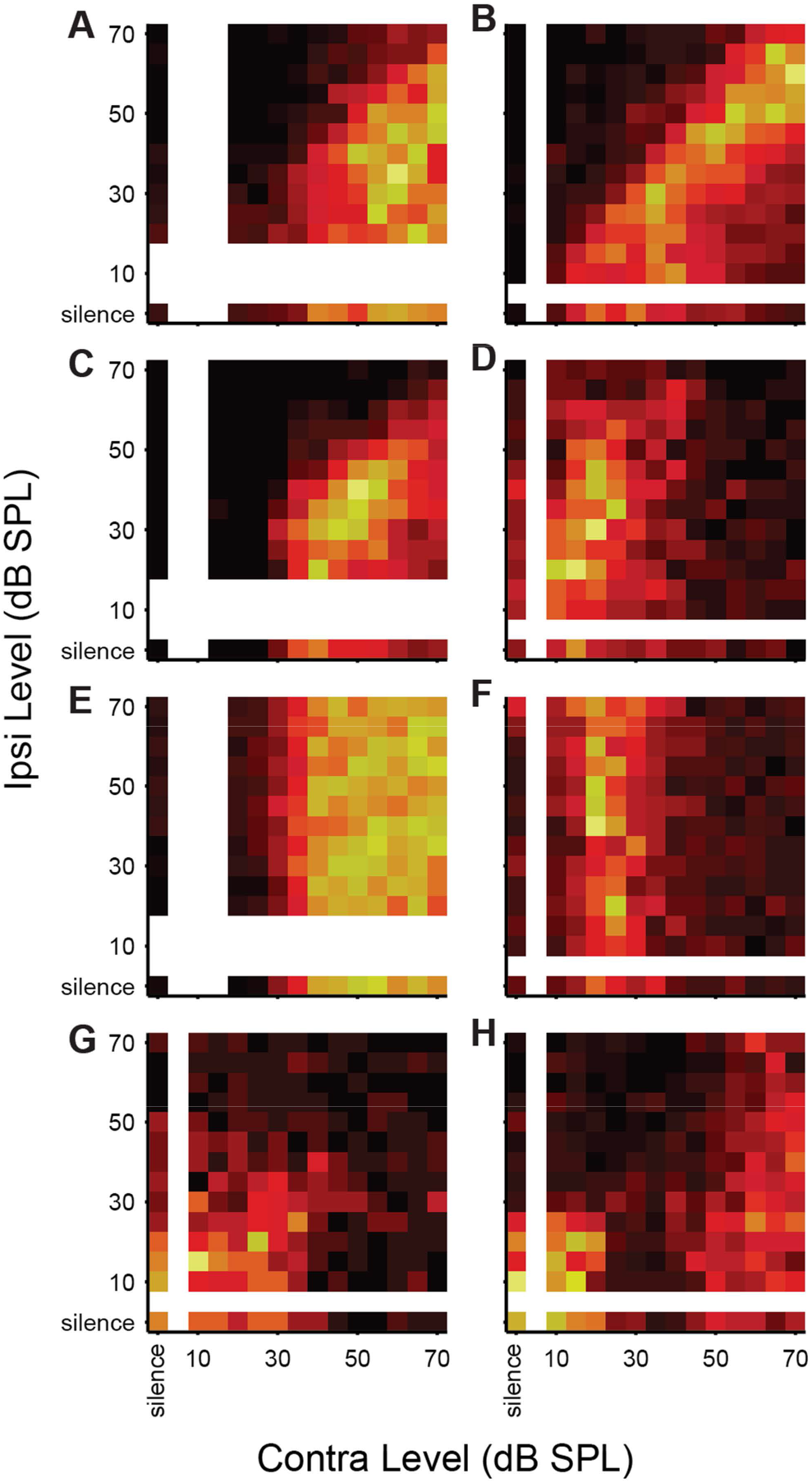
Unit types based on binaural noise response areas. **A-D**, Binaural noise response areas for 8 examplar single-units in control mice. Color scale indicates firing rate, normalized to maximum rate. **A**, Excitatory/Inhibitory (EI). **B**, Excitatory/Inhibitory (EI), contra-nonmonotonic. **C,D** Excitatory/Inhibitory with ipsilateral facilitation (EI/f), contra-nonmonotic. **E**, Excitatory/No Response (EO). **F**, Excitatory/No Response (EO), contra-nonmonotonic. **G**, Inhibitory/Inhibitory (II), **H**, Inhibitory/Inhibitory (II), contra-nonmonotonic.

